# Nicotinamide phosphoribosyltransferase activates the mitochondrial unfolded protein response to promote pulmonary arterial endothelial cell proliferation

**DOI:** 10.64898/2026.01.15.699710

**Authors:** Angelia D. Lockett, Pontian Adogamhe, Aaron Snow, Marta T. Gomes, Roberto F. Machado

**Affiliations:** Division of Pulmonary, Critical Care, Sleep and Occupational Medicine, Indiana University School of Medicine, Indianapolis, IN, USA; Department of Medicine, University of Maryland School of Medicine, Baltimore, MD

## Abstract

Vascular remodeling leading to occlusion of the pulmonary artery and increased pulmonary vascular resistance is the central feature of Pulmonary Arterial Hypertension (PAH) which leads to death due to right heart failure. The expression of Nicotinamide phosphoribosyltransferase (Nampt) is increased in the lungs and isolated PAECs of PAH patients. Inhibition of NAMPT is protective in preclinical models of pulmonary hypertension. NAMPT regulates multiple mitochondrial processes that control cell proliferation and survival. Using rodent models of PH, we demonstrated that the mitochondrial unfolded protein response (UPR^mt^) promotes vascular remodeling. Hence we, hypothesized that NAMPT activates the UPR^mt^ to promote abnormal pulmonary arterial endothelial cell proliferation. Analysis of PAECs isolated from PAH patients show increased expression of the UPR^mt^ pathway mediators, ATF-5, mtHSP70 and ClpP. Human PAEC cell lines were exposed to recombinant Nampt or transduced with lentiviral-Nampt. We observed increased phosphorylation and activation of eIF2α which permits preferential translation of the ATF-5 transcription factor. ATF-5 expression and nuclear localization was increased. Further, we observed increased expression of ATF-5 target genes, mtHSP70, HSP60, ClpP and LonP1 as well as increased PAEC proliferation. Blocking mtHSP70 function reversed the Nampt induced increase in proliferation. Neither UPR^mt^ activation nor proliferation was increased in the presence of enzymatically inactive-Nampt. Our observations demonstrate that Nampt can promote PAEC proliferation by activating the UPR^mt^ and uncovers novel potential targets to address vascular remodeling in PAH.

## Introduction

Pulmonary Arterial Hypertension (PAH) is characterized by abnormal remodeling of the pulmonary arteries of the lungs, leading to increased pulmonary vascular resistance and eventually resulting in right heart failure and death^1^. The pathobiology of PAH is multifactorial but rooted in significant cellular and metabolic dysfunction. Mitochondrial function abnormalities, including altered mitochondrial dynamics, Warburg effect and aberrant reactive oxygen species production have been centrally implicated in the development of endothelial cell dysfunction in PAH^2,3^.

Nicotinamide phosphoribosyltransferase (Nampt) is a key enzyme integral to the regulation of mitochondrial metabolic activities via NAD^+^ synthesis, ATP generation, and mitochondrial enzyme regulation^4^. Consequently, Nampt regulates fundamental biological functions, which include DNA repair, gene expression, mitochondrial function and homeostasis, cell survival, and cellular metabolism^5^. Given its role in these processes, increased Nampt expression has been observed in several tumors, likely due to the high levels of NAD^+^ required for the rapid cell proliferation^6,7^. Unsurprisingly, Nampt has been linked to several features of cancer, such as Warburg effect, tumor progression and angiogenesis^7-9^. Similarly, activation of the mitochondrial unfolded protein response (UPR^mt^) ^9^, specifically, ATF-5 and its target genes are implicated in tumorigenesis in many types of cancer^10-14^. The UPR^mt^ is an evolutionarily conserved stress-communication pathway between the mitochondria and nucleus. Activation of the ATF-5 transcriptional program creates a permissive environment for growth by trans-activating genes involved in the anti-apoptotic machinery, cell growth and migration, in addition maintaining mitochondrial homeostasis by upregulating chaperones and proteases ^15^.

We have recently demonstrated that activation of the UPR^mt^ regulates right ventricular hypertrophy and vascular remodeling in experimental PH^16^. We observed that both UPR^mt^ mediators and Nampt expression is increased in vascular cells isolated from the arteries of PAH patients and that inhibition of both pathways is protective in experimental PH ^16,17^. Given these observations and the important roles of Nampt and the UPR^mt^ in mitochondrial function, we hypothesized that Nampt initiates the activation of the UPR^mt^ to promote aberrant proliferation of PAECs.

## Methods

### Reagents, Pharmacologic Inhibitors, and Antibodies

The following antibodies were purchased from Cell Signaling Biotechnology (Danvers, MA): HSP60 (12165s), ClpP (14181s), eIF2α (2103s), PCNA (13110s), anti-mouse-HRP (7076s), anti-rabbit-HRP (7074s), and Lamin-B2-HRP (24209s). MtHSP70 is from Invitrogen (Carlsbad, CA; MA3-028). ATF-5 (NBP2-67767) and Nampt (NBP2-80036) were purchased from Novus Biological (Centennial, CO). p-eIF2α (ab32157) is from Abcam (Waltham, MA). β-Actin (A3854) was purchased from MilliporeSigma (Burlington, MA). DDK is from Origene Technologies (Rockville, MA; TA50011-100).

### Patient PAECs and hPAEC Cell lines

Approval for the use of human lung cells was granted by the Indiana University Institutional Review Board. Deidentified human pulmonary artery endothelial cells isolated from the lungs of idiopathic pulmonary hypertension IPAH, or control patients were procured from the Pulmonary Hypertension Breakthrough Initiative (Indiana University, Indianapolis, IN). Primary hPAEC cell lines were purchased from Lonza (Walkersville, MA; CC-2581) or ATCC (Manassas, VA; PCS-100-023). Cells were cultured at 37°C in Endothelial Cell Growth Medium 2 (EGM2, Lonza, CC-3162) and were used between passage 4 to 8 for all studies.

### Recombinant Nampt treatment and lentiviral transduction

hPAECs were exposed to recombinant (MBL Life Science, Carlsbad, CA; CY-E1251) for 24h or vehicle control (20mM Hepes-KOh, pH 7.5, 1mM DTT, 50mM NaCl, 50% glycerol). Lentiviral Nampt-DDK, Nampt-dominant negative (Nampt-DN, H247A) and control vectors were designed, packaged and purchased from Vector Builder Inc (Chicago, IL). Human Nampt was cloned into a lentiviral plasmid that expressed hNampt-DDK (VB220309-2189 wac) or hNampt-DN (VB220830-1074 gtp) downstream of the EF1A promoter and EGFP downstream of the CMV promoter. hPAECs (300,000 cells/well) were seeded onto 6-well plates and allowed to attach overnight in a tissue culture incubator. Lentiviral particles were diluted in EGM2 with sufficient viral titer to achieve MOI 20 and then added to cells. After 24h the lentiviral containing media was replaced with EGM2 and left for 48h followed by downstream applications.

### Whole cell extraction, nuclear extraction, and Western blotting

For analysis of whole cell extracts, cells were rinsed with cold PBS, collected in 1x cell lysis buffer (Cell Signaling; 9803) containing 1x protease/phosphatase inhibitor cocktail (ThermoFisher; 78442), incubated on ice for 15 min and centrifuged at 14,000 x g for 5 min to collect protein. For nuclear extractions, cells were rinsed with cold PBS and centrifuged at 500 x g for 5 min in 1mL of cold PBS. Cell pellets were gently resuspended in nuclear extraction buffer (10mM Tris-HCl pH 8.0, 140mM NaCl, 1.5mM MgCl2, 0.5% NP-40 (v/v), 1x protease/phosphatase inhibitor cocktail) followed by incubation on ice for 5 min. The cytoplasmic supernatant was removed prion to suspending the nuclear pellet in cell lysis buffer and incubating on ice for 30 min followed by centrifugation at 14,000 x g for 5 min. Proteins were quantified and resolved by SDS-PAGE and transferred to PVDF membranes. Western Blotting was performed by incubating with primary antibodies (1:1000, overnight, 4◦ Celsius). HRP conjugated secondary antibodies were used at 1:5000 for 1h at room temperature.

### Statistical Analysis

Statistical analysis of experimental data was performed using GraphPad Prism 5.1 (GraphPad Software, Inc., La Jolla, CA). Results are expressed as mean ± SEM from at least three experiments. Student *t* test and analysis of variance were used to compare two and three groups, respectively. *P* less than 0.05 was considered statistically significant.

## Results

### The UPR^mt^ is activated by Nampt in PAECs

We have shown that the UPR^mt^ contributes to the PH pathogenesis, as pharmacological inhibition of mtHSP70 is protective against both hemodynamic changes and vascular remodeling in experimental PH models^16^. Our previous studies also demonstrate that heterozygous genetic deletion of Nampt is sufficient to mitigate increases in right ventricular systolic pressure (RVSP) as well as RV hypertrophy and vascular remodeling, demonstrating a protective effect in rodent models of PH. We further demonstrated that Nampt is upregulated in the lungs and PAECs of idiopathic PAH patients^17^. Hence, we assessed the expression level of UPR^mt^ proteins in PAECs isolated from IPAH patients. Compared to non-disease controls, there was a significant increase in the levels of ATF-5, mtHSP70 and ClpP in patient cells (**Figure 1**). The integrated stress response leads to eIF2α phosphorylation and activation and phosphorylated eIF2α promotes ATF-5 translation and propagation of the UPR^mt^. Hence, we assessed the effect of Nampt on eIF2α activation and ATF-5 expression and localization by exposing human pulmonary artery endothelial cells (PAECs) to recombinant Nampt (rNampt) or by overexpressing Nampt (L-Nampt) using lentiviral delivery. Nampt led to increased phosphorylation of eIF2α (**Figure 2A**) as well as to increased ATF-5 expression (**Figure 2B**) and nuclear translocation (**Figure 2C**). Assessment of the expression level of ATF-5 target genes, demonstrate that both rNampt and L-Nampt led to significantly increased expression of mtHSP70, HSP60, ClpP and LonP1 as well as the proliferation marker, PCNA (**Figure 3**).

**Figure 1.**
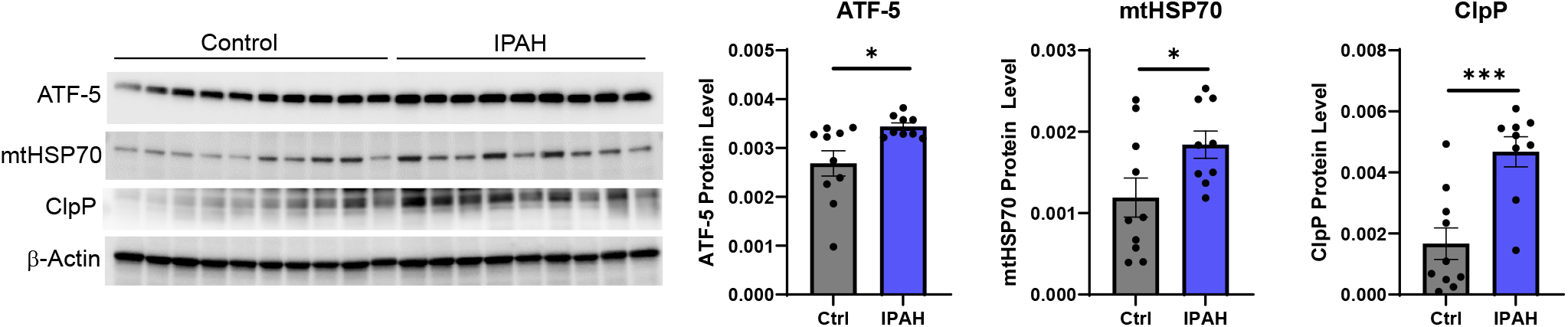
The expression of UPR^mt^ mediators is increased in PAECs isolated from IPAH patients. Whole cell extracts were collected to assess the expression of UPR^**mt**^ proteins. Western blot and densitometry demonstrate elevated levels of ATF-5, mtHSP70 and ClpP. Results are expressed as mean + SEM versus control; *p<.05, ***p<.0008

**Figure 2.**
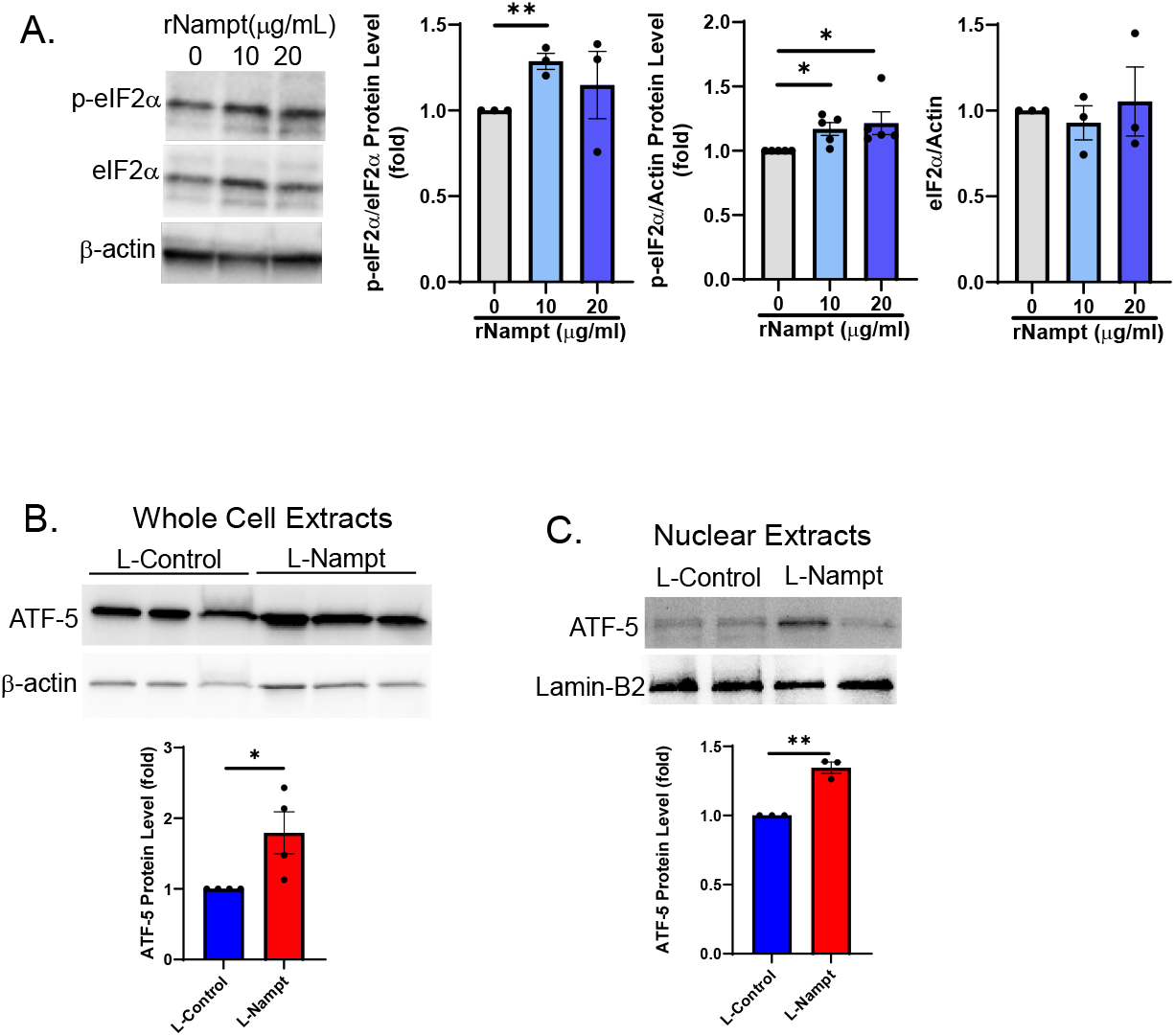
Nampt promotes activation of the UPR^mt^ in hPAECs. Western blot and densitometry show that recombinant Nampt (A; rNampt 24h) led to increased eIF2α activation (p-eIF2α), and that lentiviral overexpression of DDK-tagged Nampt (L-Nampt; MOI 20, 48h) lead to increased whole cell (B) and nuclear levels of ATF-5 (C). Results are expressed as mean + SEM; *p<.05, **p<.006

**Figure 3.**
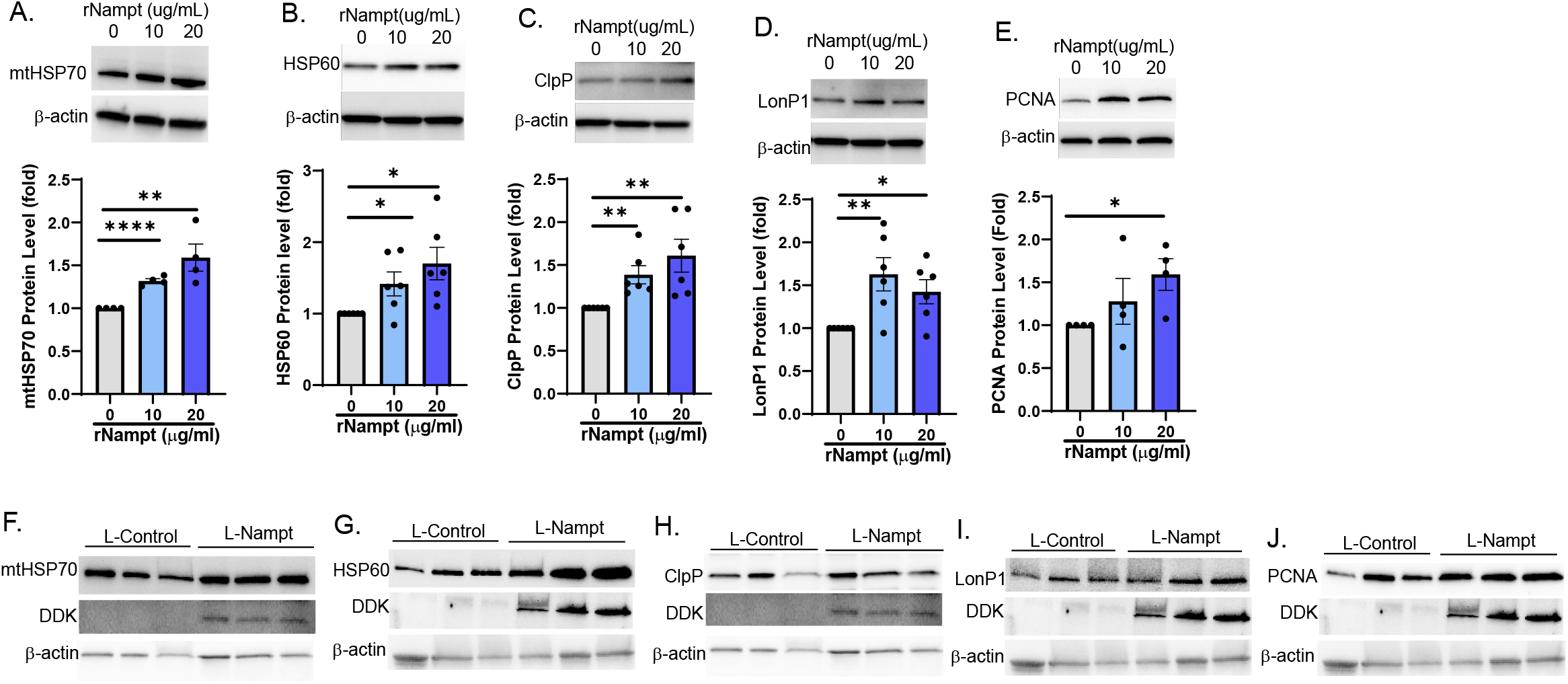
Nampt leads to upregulation of ATF-5 inducible genes in hPAECs. Western blot and densitometry show that recombinant Nampt (A-D; rNampt 24h) and lentiviral overexpression of DDK-tagged Nampt (F-I; L-Nampt; MOI 20, 48h) led to increased whole cell levels of mtHSP70, HSP60, ClpP and LonP1 and increased proliferation (E,J). Results are expressed as mean + SEM; *p<.05, **p<.009, **** p<.0001

### The enzymatic activity of Nampt is required for UPR^mt^ activation and PAEC proliferation

The enzymatic activity of Nampt catalyzes the rate-limiting step of NAD^+^ synthesis from nicotinamide. NAD^+^ is an essential metabolite in glycolysis, and it is consumed to promote glycolysis^18,19^. In PAH, increased glycolysis is a well-established pathological feature that promotes increased proliferation and induces vascular remodeling^20-23^. Noteworthy, low NAD^+^ levels are associated with UPR^mt^ activation and increased intestinal aging ^24^. Hence, we determined if the enzymatic activity of Nampt is necessary for activation of the UPR^mt^ by overexpressing a dominant negative mutant (Nampt-DN) with a point mutation (H247A) in the catalytic domain. While Nampt overexpression led to increased expression of UPR^mt^ mediators and promoted proliferation, there was no significant difference between Nampt-DN and control transduced cells (**Figure 4**). These data suggest that the NAD^+^ generating activity of Nampt is involved in the UPR^mt^ mediated proliferation of PAECs.

**Figure 4.**
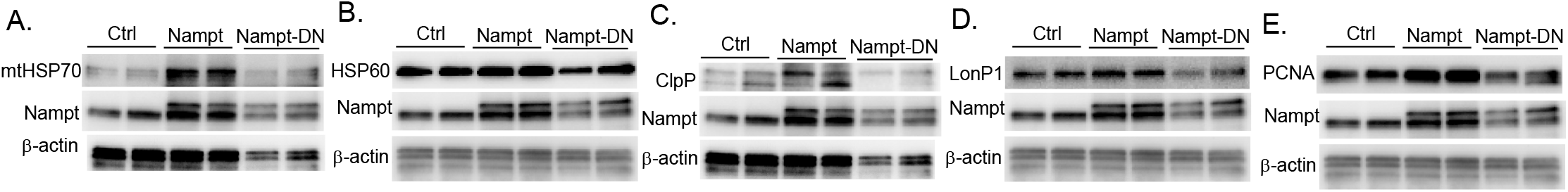
Nampt enzymatic activity is necessary for UPR^mt^ activation. Western blot and densitometry showing reduced expression of UPR^mt^ proteins in hPAECs transduced with enzymatically inactive Nampt (MOI 20, 48h) comparted to wildtype Nampt (MOI 20, 48h) or to controls. Results are expressed as mean + SEM; *p<.05, **p<.003, *** p<.0008

### Inhibition of mtHSP70 prevents Nampt induced proliferation of PAECs

We previously established that Nampt has a profound effect on vascular remodeling in PH in that it induces proliferation of both PAECs and PASMCs. Furthermore, conditioned media from PAECs promotes PASMC proliferation, presumably via paracrine signaling of Nampt since a Nampt inhibitor, FK866, blocks the proliferation^17^. We tested whether the mechanism by which Nampt promotes PAEC proliferation involves UPR^mt^ signaling. We confirmed that Nampt promotes PAEC proliferation by assessing the protein level of PCNA and found that concurrent with UPR^mt^ activation, there was an increase in PCNA expression (**Figure 3E, J**). Further, the enzymatic activity of Nampt was necessary to promote proliferation (**Figure 4E**) . To examine the role of the UPR^mt^ in Nampt induced proliferation, we overexpressed Nampt for 48h which is sufficient to induce proliferation as shown in figure 3. We then blocked mtHSP70 signaling for 24h using an allosteric pharmacological agent, Mkt-077, which crosses the mitochondrial membrane to specifically bind mtHSP70 ^21^. Blocking mtHSP70 reversed the Nampt-induced increase in PAEC proliferation as PCNA expression was decreased (**Figure 5**). mtHSP70 complexes with CoxIV isoform 2, an isoform that is predominantly expressed in the lung, to promote assembly of cytochrome c oxidase in the electron transport chain. Hence, CoxIV protein levels correlate with the ability of mtHSP70 to bind its targets^21^. Our analysis of total CoxIV levels indicate that Nampt promoted CoxIV expression and demonstrated that Mkt-077 blocked mtHSP70 binding activity as CoxIV expression was decreased in Nampt overexpressing cells exposed to Mkt-077 (**Figure 5**). Furthermore, mtHSP70 expression was also decreased. Taken together, these data indicate that Nampt modulates mitochondrial signaling by promoting activation of the UPR^mt^ to induce PAEC proliferation.

**Figure 5.**
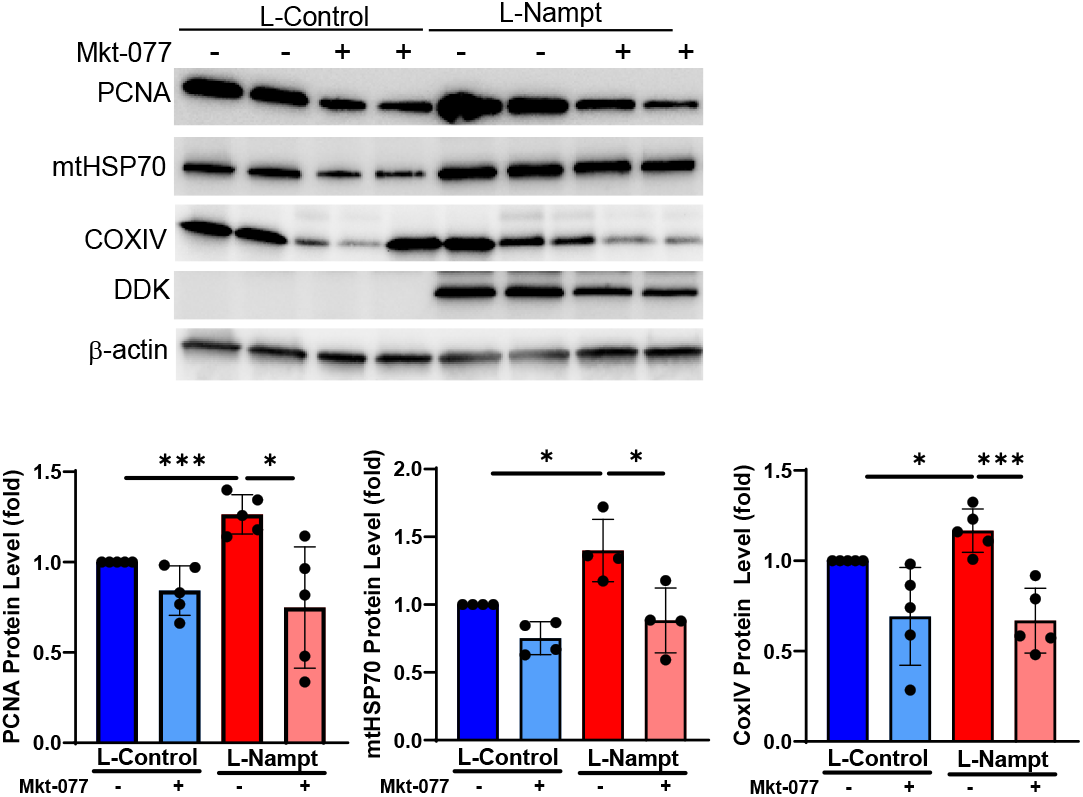
Inhibition of mtHSP70 mitigates Nampt-induced hPAEC proliferation. L-Nampt was overexpressed (MOI 20, 48h) prior to treatment with Mkt-077 (1μM). Western blot and densitometry demonstrating decreased PCNA, mtHSP70 and CoxIV expression in Nampt over expressing cells treated with Mkt-077 (24h). Results are expressed as mean + SEM; *p<.05, **p<.006, *** p<.0005

## Discussion

Nampt plays an important role in mitochondrial function by maintaining the supply of NAD^+^, an essential metabolite that is needed for energy homeostasis and the regulation of NAD^+^ dependent enzymes^7,9^. These enzymes are critical to regulating cellular and mitochondrial homeostatic functions such as apoptosis, mitochondria biogenesis, DNA repair, redox balance and cellular stress regulation . We previously reported that Nampt expression is increased in the lungs and pulmonary arterial endothelial cells of IPAH patients and that Nampt secreted from PAECs may mediate vascular cell crosstalk as conditioned media from IPAH patient PAECs promotes PASMC proliferation^17^. Both in vivo and in vitro analyses demonstrated that Nampt promoted pulmonary vascular remodeling and that inhibition of Nampt either via genetic deletion or pharmacological inhibition was protective, indicating its potential as a therapeutic target in PAH^17^. Dysregulation of multiple mitochondrial processes, i.e. respiration, ROS, glycolysis and fission, are key features of PAH that contribute to vascular remodeling. We present herein a novel association between activation of the mitochondrial unfolded protein stress response and Nampt regulation of PAEC growth. Our studies demonstrate that overexpression of Nampt in PAECs results in increased UPR^mt^ activation as well as increased PAEC proliferation and that these functions of Nampt are dependent on its enzymatic activity.

The UPR^mt^ is an evolutionarily conserved stress response that regulates mitochondrial health and survival under stressful conditions. It has been studied extensively in *Caenorhabditis elegans* where it is regulated by the activation transcription factor associated with stress 1 (ATFS-1). Under normal conditions, ATFS-1 is imported from the cytoplasm into mitochondria where it undergoes degradation. However, under stressful conditions that impact the ability of mitochondria to import proteins, ATFS1 translocates to the nucleus to transactivate chaperones and proteases that mediate the UPR^mt^ . This mirrors the function of the mammalian homologue, ATF-5, which induces the expression of chaperones (i.e. mtHSP70 and HSP60) and proteases (ClpP an LonP1) to regulate mitochondrial homeostasis. Our assessments of PAECs isolated from IPAH patients revealed increased expression of UPR^mt^ genes; ATF5, mtHSP70, and ClpP. Furthermore, when we assessed the role of Nampt on promoting upregulation of the UPR^mt^, we observed increased expression and nuclear localization of ATF-5 which was commensurate with increased phosphorylation and activation of eIF2α. Each ATF-5 inducible protease (ClpP and LonP1) and chaperone (mtHSP70 and ClpP) that we assessed was also increased.

Prior mechanistic studies revealed that the UPR^mt^ supports a pro-survival, anti-apoptotic, pro-proliferative phenotype^16,36,38,39^. Hence, while UPR^mt^ activation is beneficial for improving mitochondrial health and function in response to acute stress, prolonged activation can be detrimental by allowing for the survival and proliferation of dysfunctional cells, which has been shown to promote tumor growth and progression^15,36,40-42^. For example, during mitochondrial stress the UPR^mt^ promotes upregulation of metabolic genes which promote glycolysis while simultaneously limiting transcription of TCA and oxidative phosphorylation genes^37,43^. Although, these alterations maintain cellular function short term, they pose a significant long term risk as these alterations are characteristic of highly proliferative cells as occurs in PAH^44^. This is in line with our observation that Nampt induced proliferation of PAECs occurs via activation of the UPR^mt^. Of note, overexpression of mtHSP70, in particular, can lead to loss-of-function of the p53 tumor suppressor^45^. The proliferative, tumorigenic function of mtHSP70 led to cancer clinical trials in which Mkt-077 was used to selectively inhibit the chaperone activity of mtHSP70^46^. The mtHSP70 target, CoxIV isoform 2, is induced by hypoxia to promote angiogenesis and tumorigenesis ^47^. Furthermore, a key mediator of PAH, HIF1α, also regulates the expression of CoxIV during low oxygen conditions to meet cellular energetic demands; hence CoxIV isoform 2 serves as an oxygen sensor. ^48,49^. We observed that mtHSP70 inhibition attenuated the Nampt mediated increase in PAEC proliferation and CoxIV expression. The effect of Nampt and mtHSP70 inhibition on CoxIV and PCNA expression corroborates the role of the UPR^mt^ in mediating Nampt induced PAEC proliferation.

Our findings provide valuable insight into the mechanisms involved in Nampt regulation of the pulmonary vascular endothelial cell growth that contributes to vascular remodeling in PAH. The discovery that the UPR^mt^ acts downstream of Nampt presents novel targets to investigate as potential points of therapeutic intervention. Future studies are focused on both investigating the effect of NAD/NADH on UPR^mt^ activation in the pulmonary vasculature as well as validating these findings in relevant *in vivo* models of experimental PAH.

## Funding Sources

This work was supported by grants from the National Heart, Lung and Blood Institute of the National Institutes of Health (NHLBI-K01HL164874, NHLBI-1R01HL158-01A1 and the Indiana University Center for Translational Science-Project Development Team. ALA - Award ID 1278310, CTSI Indiana-BRG (Biomedical Research Grant).

## Notes

**Disclosures** We have no known conflicts of interest to disclose.

### Competing Interest Statement

The authors have declared no competing interest.

